# Development of Tribo-Electroceutical Fabrics for Potential Application in Self-sanitizing Personal Protective Equipment (PPE)

**DOI:** 10.1101/2021.02.23.432624

**Authors:** Sayan Bayan, Aniruddha Adhikari, Uttam Pal, Ria Ghosh, Susmita Mondal, Soumendra Darbar, Tanusri Saha-Dasgupta, Samit Kumar Ray, Samir Kumar Pal

## Abstract

Attachment of microbial bodies including coronavirus on the surface of personal protective equipment (PPE) is found to be potential threat of spreading infection. Here, we report the development of a novel tribo-electroceutical fabric (TECF) consisting of commonly available materials namely Nylon, and Silicone Rubber (SR) for the fabrication of protective gloves on Nitrile platform, as a model wearable PPE. A small triboelectric device (2 cm × 2 cm) consisting of SR and Nylon on Nitrile can generate more than 20 volt transient or 41 µW output power, which is capable of charging a capacitor up to 65 V in only ∼50 sec. The novelty of the present work relies on the TECF led anti-microbial activity through the generation of an electric current in saline water. The fabrication of TECF based functional prototype gloves can generate hypochlorite ions through the formation of electrolysed water upon rubbing them with saline water. Further a computational modelling has been employed to reveal the optimum structure and mechanistic pathway of anti-microbial hypochlorite generation. Detailed anti-microbial assays have been performed to establish effectiveness of such TECF based gloves to reduce the risk from life threatening pathogen spreading. The present work provides the rationale to consider the studied TECF, or other material with comparable properties, as material of choice for the development of self-sanitizing PPE in the fight against microbial infections including COVID-19.

## Introduction

The respiratory diseases such as COVID-19 can spread from the virus-laden respiratory droplets emerging out from the coughing or sneezing of any infected person.^1^ In practice, health workers are continuously exposed to such life threatening biological agents and can become potential carriers of the diseases. To minimize the life risk of the health workers the use of Personal Protective Equipments (PPEs) in the form of gowns, face masks, gloves, goggles, face shields etc. are highly recommended.^2^ The proper use of PPEs is considered as a common precautionary measure to reduce the risk of contamination from blood or other body fluids of patients. However, during the COVID-19 pandemic, the whole world has experienced an unprecedented challenge in PPE supply for active patients as well as health workers.^3-5^ Consequently it has been realized that the long duration use of PPEs has led to the adverse effect due to the negligence of hand hygiene practices and recycling following the disease protocols in regular intervals. For disinfection and reuse of PPEs a number of routes such as exposure to heat (autoclave), ultraviolet ray-C, alcohol, washers etc. have been recommended.^5^ However, most of these methods are time consuming and face affordability issues by economically weaker sections of the society. So, the development of self sanitizing PPEs can be a novel and alternate solution for the all sections of the society including the health workers.

The contact electrification process (also known as tribo-electrification) has attracted the attention of medical research community for healthcare application.^6-8^ The tribo-electrification involving the phenomena of tribology with subsequent charge transfer/exchange has led to the development of energy harvesting devices called triboelectric nanogenerators (TENG).^9-13^ In this context, a TENG can be exploited as a health monitoring sensor as well as power generator, where the electrical signals in response to a stimuli originating from human body are able to power up energy storage devices that can operate the sensor.^6, 14-18^ For example, activities like muscular motion or abdominal respiration, where the body displacement is significant, can be easily detected by TENG^7, 8^ and can power the sensing systems to detect heart beat, body temperature, and lactate concentration etc.^14-16^ The potential applicability of TENG in antibacterial wearable systems has also been reported.^19-21^ The fabrication of self-powered filter by combining TENG and photocatalysis technique has been found to be efficient to protect from harmful organic vapour pollutants, which are believed to be the cause of diseases like asthma and chronic bronchitis.^19^ On the other hand, antibacterial composite film-based TENG has been demonstrated to be useful to protect athlete’s foot from fungal infection effectively.^20^ Excellent antibacterial activity along with capability of removing fine particle (PM2.5) by the virtue of cellulose fiber based TENG has also been reported.^21^

Here, we first report the development of a tribo-electroceutical fabric (TECF) for the fabrication of personal protective equipment like protective gloves, as a representative wearable PPE against deadly viruses in a situation like COVID 19 pandemic. We demonstrate the tribo-electroceutical fabric (TECF) based gloves deploying the virtue of contact electrification using commonly available materials like nylon, teflon or silicone rubber (SR). The TECF based on nylon or SR, act as contacting materials on nitrile platform, a common material for hand gloves, and can generate electrical power. The developed TECF based gloves when rubbed in presence of saline water produce hypochlorous acid or hypochlorite ions, which lead to their self-sanitization while using for a longer duration. The production of the hypochlorite ions has been characterized *in vitro* through a redox dye degradation as well as colorimetric test. In addition, theoretical modelling has been performed to establish the concept of electrolysis and the consequent formation of hypochlorous acid. Finally, the efficacy of the TECF has been revealed by demonstrating the sterilizing effect through killing a vast majority of *Pseudomonous auregionosa* bacteria, which has similarities with SARS CoV-2 virus. The present investigation suggests that TECF based gloves can effectively reduce the chance of contamination from microorganisms that are spread by direct or indirect contact through a self-sanitization process, making them attractive to prevent further spread of the diseases from an otherwise uninfected personnel.

## Experimental and Computational Details

### 1. Development of triboelectric materials

Nylon layer, one of the tribo-eletroceutical surfaces in the present work, was prepared using commercially available nylon beads (∼ 0.04 gm), which were mixed in ∼ 10 ml acetone solution and stirred at ∼ 400 rpm for 15 min by maintaining a temperature of 50 °C. The resulting viscous solution was drop casted on commercial nitrile gloves and was allowed to dry in a hot air oven. Silicon rubber (SR) layer, acting as the counter tribo-material, was prepared from liquid SR and a catalyst (e.g. dibutyl tin dilaurate), which were mixed in 13:1 ratio and casted on nitrile gloves. The desired prototype of nylon and SR coated nitrile gloves were obtained upon curing the samples in a hot air oven at ∼ 70 °C.

### 2. Exploration of triboelectric properties

For investigating the triboelectric characteristics, approximately 2 cm × 2 cm nylon and SR coated nitrile sheets were attached to the adhesive aluminium tape. An adhesive aluminium foil was used to make the back contact of the triboelectric layers. An indigenously developed system with two wooden slabs separated by four springs at the corners was deployed to measure the triboelectric characteristics under tapping conditions. The triboelectric layers were attached to the two inner faces of the system so that the layers could come in contact with each other under pressed condition. Further for sliding/rubbing mode, the triboelectric layers were fixed to a polyethylene terephthalate (PET) platform with a spacer such that the horizontal movement can be executed.

### 3. Characterization

The electrical properties of the nanogenerators were studied using a digital oscilloscope (Scientific) and Keithley 2450 source meter. Optical absorbance was recorded using an UV-Visible spectrophotometer (Jasco) at room temperature.

For dye degradation experiment, the TECF based gloves were submerged in methylene blue solution (4.5 μM) in normal saline (0.9% NaCl) condition. The system was continuously rubbed and the absorbance was measured from aliquots collected from the sample at 5 min time intervals.

The bacterial strains of gram-negative *Pseudomonous auregionosa* used in this study was obtained from Dey’s Medical (Kolkata, India), while *Escherichia coli (E. coli)* (CGMCC 1.8723) was obtained from the Department of Biochemistry, University of Calcutta, India. The glass wares, suction nozzles, and culture medium were sterilized in an autoclave at a high pressure of 0.1 MPa and a temperature of 120 °C for 30 min before experiments. Bacteria cultures were cultivated in sterilized Luria-Bertani (LB) broth and incubation at 37 °C with a shaking incubator for 24 hours.

The colony count method or the kinetic test was used to estimate antibacterial properties through the concentration of the survival colonies bacteria in co-cultured solution. First, original bacterial suspensions were washed three times with phosphate-buffered saline (PBS; pH 7.4) solution to a concentration of 10^8^ CFU/ml. At the second step, they were poured onto nylon-nylon (TECF Control) or SR-nylon (TECF) layers and continuously rubbed for 25 mins. At third step, the rubbed bacterial solution was diluted five times to a certain concentration. The resulting bacterial PBS suspensions (100 μL) were spread on gelatinous LB agar plates, and cultured at 37°C for 24 hours. The number of survival colonies was counted manually. For growth curve analysis, the five times diluted bacterial solution was added to LB and kept at 37°C. The absorbance at 660 nm was monitored at a regular interval (1 h) to assess the growth kinetics.

The bacteria cells after treatment with TECF, were stained with DAPI and PI. The DAPI can stain all cells while the PI can only stain the living cells, and thus the red/blue ratio was obtained to assess the survivality of *E. coli*.

### 4. Computational study

The fabricated TECF was simulated using the electrostatic module of the COMSOL Multiphysics simulator. The potential difference (*U*) was obtained by solving the equation: *U = σ∆d/ε*_0_, where *σ* is the surface charges density; *ε*_*0*_ is the vacuum permittivity and *∆d* represents the gap distance between the two tribo-materials with electrodes.^22^ In the simulations, a charge density of −10 nC/m^2^ was assumed on the SR layer surface. The bottom electrode was taken as a reference in the simulation and was grounded for the purpose.^23,24^

The secondary current distribution through the electrolyte is simulated using the electrochemistry module of COMSOL. Electrolyte in between the two triboelectric materials can be modeled as an electrolysis cell with an anode and a cathode. Chlorine/chloride reduction (E_eq,Cl_) potential of +1.23V was applied to the anode where the chlorine gas evolution takes place. Water/hydrogen reduction potential (E_eq,H_) of -0.828V was applied to the cathode where reduction of water to hydrogen takes place releasing also hydroxide ions. Therefore, the thermodynamically equilibrium cell potential becomes, E_eq,cell_ = E_eq,Cl_-E_eq,H_ = +2.058V. However, at this potential, reaction does not take place due to some kinetic loss. The chloride oxidation reaction at anode is very fast (reversible reaction) and departs very little from the equilibrium potential. On the other hand, hydrogen evolution reaction is inherently slow (irreversible reaction and the cathode material not being a perfect catalyst results in a kinetic limitation) and requires a potential, E_pol_, significantly greater in magnitude than the equilibrium potential (E_eq,cell_) to achieve a reasonable current density to make up for the kinetic losses and drive the reaction. This potential is called the overpotential and the electrode is said to be polarized. Here, the cell is polarized to +10V vs the cell equilibrium to drive the reaction. Thus, the overall cell potential or the electrode to electrode potential becomes E_cell_ = E_eq,cell_ + E_pol_ = +12.058V. Common Butler-Volmer equation ^25^ was used to parameterize the slow hydrogen evolution reaction at cathode with an exchange current density of 1mA/m^2^ on the reacting surfaces. Electrolyte conductivity of the free electrolyte is defined as 5 S/m. A stationary study was performed to describe the steady state current in the system.

Quantum chemical study of Cl^2^ splitting and HOCl formation was carried out with B3LYP exchange correlation functional ^26^ and 6-31g(d) basis set on all atoms as implemented in Gaussian 16 software.^27^ Bond distance between two chlorine atoms was perturbed to obtain the potential energy diagram in presence of explicit water molecules.

Structure and sequence of the cell surface adhesion lectin protein lecA of Pseudomonous auregionosa was obtained from Protein Data Bank (PDB ID: 4YWA). Structure and sequence of the receptor binding domain (RBD) of SARS-CoV-2 was also obtained from the PDB (ID: 6LZG). Sequence alignment was performed using Clustal Omega web server.^28^ Similar amino acids in the binding sites of the two proteins were mapped using PyMOL molecular graphics software.

## Results and Discussion

Initially the triboelectrical output of the TECF pairs has been investigated using a indigenously built spring based experimental arrangement. Figure 1(a) shows the schematic representation of the experimental set-up used for measuring the electrical output of triboelectric devices. The mechanical pressing and releasing actions of the spring based system lead to the contact and separation states of two layers. Initially, the tribo-electrification has been investigated in three TECF pairs: nitrile-nylon, teflon-nylon and SR-nylon. The active materials have been chosen based on their affinity towards positive and negative charges following triboelectric series. As shown in Figure 1 (b) for all the three pairs, recurring negative/positive pulses of open circuit voltage are observed upon repetitive pressing/releasing actions (an enlarged view of nitrile-nylon system presented in the inset). The Nitrile-Nylon TECF pair exhibits the lowest output voltage, while the SR-Nylon pair showing the highest one. The SR-nylon pair displays a maximum open circuit voltage of ∼ -20 V during the pressing action. The output generation by the tribo-system has been verified by switching the polarity of the measuring instrument (Figure S1 in supporting information). The origin of output voltage can be explained by the well established theory based on the triboelectric charge generation and subsequent potential development, shown schematically in the supporting information (Figure S2). On the other hand, the variation in output voltage for the three pairs is attributed to the varying electron accepting or donating tendency, as predicted by the triboelectric series. The power generation has been tested by varying the load resistance (ranging from 45 kΩ to 15 MΩ) and the SR-nylon device displays a maximum power of ∼ 41 µW/cm^2^ at 200 kΩ, as presented in Figure S3. The voltage generated by the contact electrification has been used to charge a commercial capacitor (0.26 μF), as shown in Figure 1(c) using a rectifier circuit (shown in the inset). The repetitive pressing leads to the charging of the capacitor with a development of ∼ 65 and 74 V, respectively, for teflon and SR based pairs within ∼ 50 s. Multiple commercial LEDs can be powered with bright intensity using the charged capacitor, as shown in the inset of Figure 1(c).

**Figure 1:**
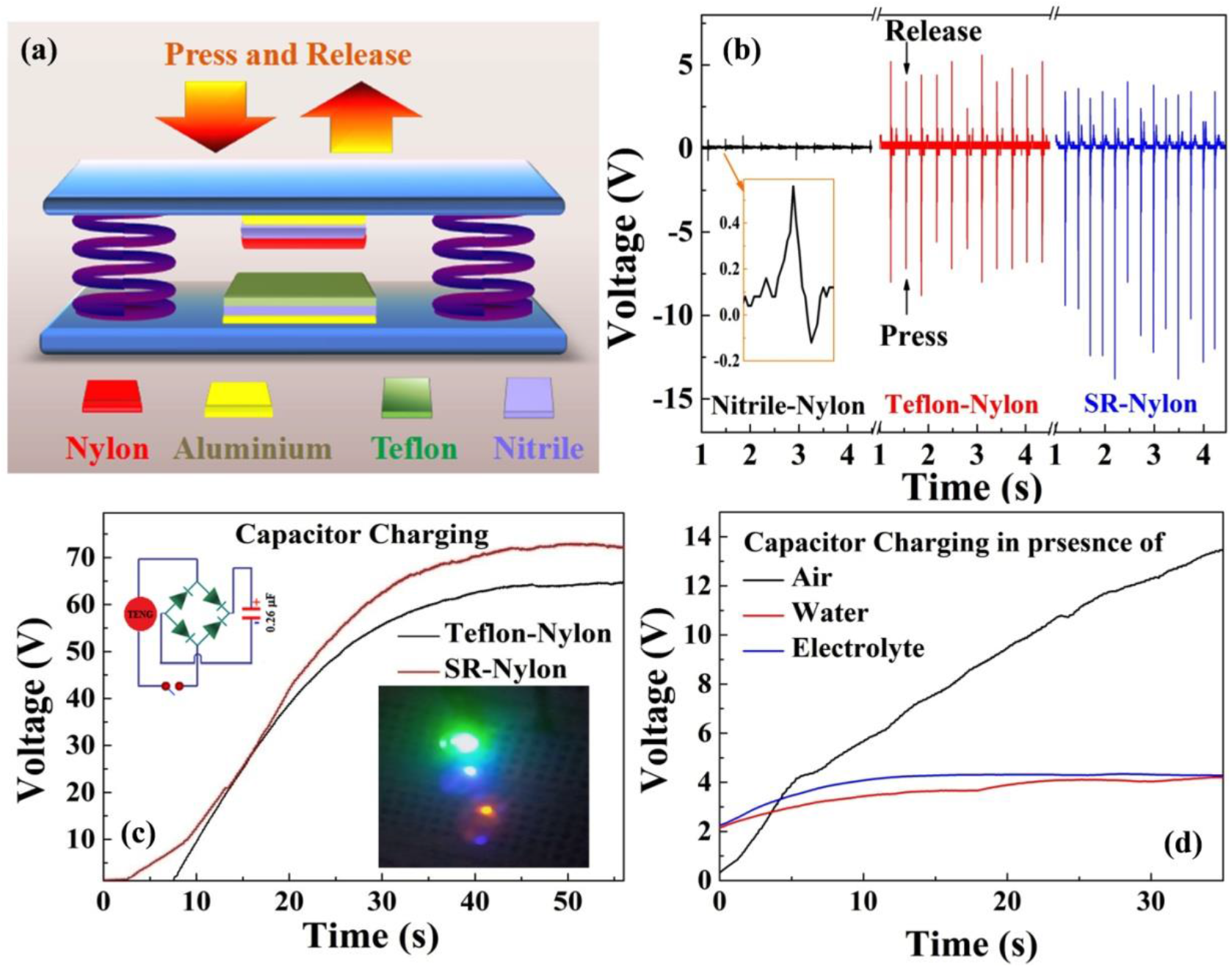
(a) Scheme of the tribo-device for contact electrification measurements, (b) Output voltage pulses upon press-release action with the inset showing the output for nitrile-nylon pair, (c) Charging of a capacitor (0.26 µF) by press-release action of teflon-nylon and SR-nylon pairs, while the inset showing the scheme of a bridge rectifier circuit and intense glow of multiple commercial LEDs through the charged capacitor. (d) Capacitor charging by rubbing in presence of different mediums.

With an aim to deploy the TECF in wearable kits, the performance of the two triboelectric materials has been checked upon rubbing/sliding action. The output voltage under different rubbing arrangements of the contact layers are presented in Figure S4(a,b), while the charging profile of a capacitor under rubbing actions with different frequencies is shown in Figure S4(c). Next, to generate a short circuit current between the tribo-layers, the contact-separation experiments have been performed in sliding mode with an intermediate medium between the layers. The results of the sliding experiments in presence of deionized (DI) water and sodium chloride (NaCl) electrolyte solution are presented in Figure 1(d). In contrast to air medium, the voltage developed across the capacitor is quite low when the sliding experiments are carried out in presence of DI water and NaCl solution. In both situations, there is a signature of saturation. It indicates that the liquid medium present within the two dielectrics, experiences two opposite type of charges at the two liquid/dielectric interfaces and thus allows substantial electron flow leading to a situation of internal short circuit condition. However, owing to the finite resistance of water or electrolyte solution, the potential doesn’t drop to zero and thus a saturated value of voltage is maintained across the capacitor.

To visualize the contact electrification process in the presence of varying medium (dielectric) between the tribo-layers, finite element simulation has been carried out using COMSOL Multiphysics software. Figure 2(a,b) shows the potential distribution between the two electrodes in air medium with a varying separation of 1.00 and 10.0 mm. It is observed that a maximum potential difference of ∼ 18 V can be achieved when the electrode separation reaches 10 mm. This is because, as the two layers are separated, mechanical work has to be done against the electric field and thus results in the increase in potential difference between the top and bottom electrodes. On the other hand, the potential distribution becomes negligible, under the same condition for the two triboelectric materials when the intervening medium is replaced from air to water or an electrolyte solution (Figure 2(c)). These calculated results are in agreement with our observed experimental results, with the output voltage diminishing when the contact electrification experiment is performed in presence of water or electrolyte medium, as shown in Figure 1(d). It is expected that the internal short circuit leads to the electrolysis of the electrolyte solution, where the oxidation of chloride to chlorine takes place at the anode. On the other hand, the reduction of water into hydrogen gas takes place at the cathode along with the release of hydroxide ion. The solution of the secondary current distribution calculation (Figure 2(d)) represents the electrolyte potential in the cell and reveals a higher electrolyte potential in the anode side of the cell. The arrow plot indicates the direction of the current flowing from anode to cathode. Electrolyte to electrode potential difference at the cathode is 1.72 V, which corresponds to the kinetic loss of the potential drop. Owing to this kinetic loss in the system, a large over-potential is required to drive the reaction. Nevertheless, the evolution of the circulating current in TECF is quite attractive for achieving self-sanitation effect, which will be discussed later.

**Figure 2:**
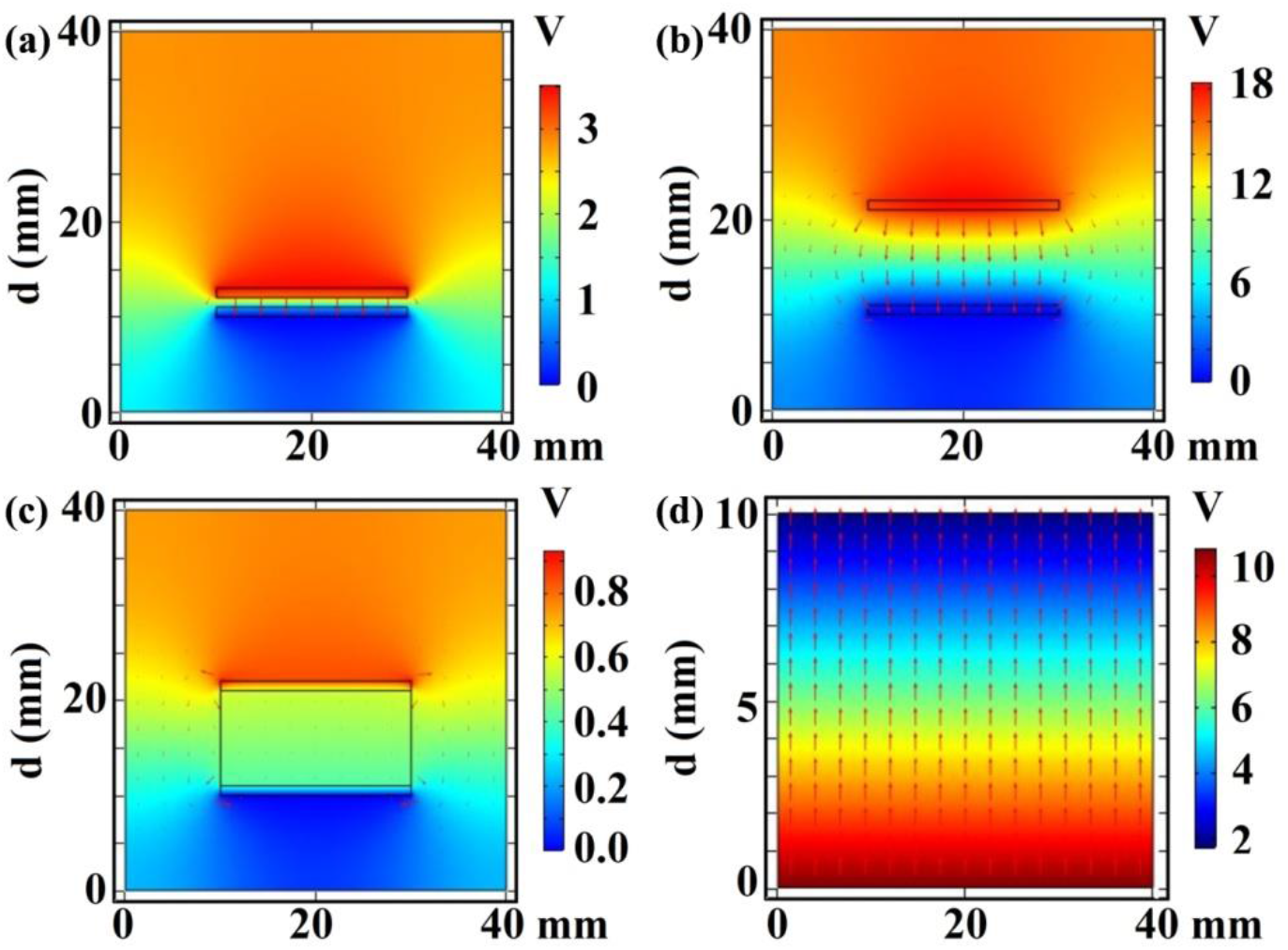
Finite element simulation of potential distribution in SR-Nylon TECF with electrode separation of (a) 1.0 mm and (b) 10.0 mm. (c) Potential distribution in presence of water/saline water between dielectrics with electrode separation of 10 mm. (d) Secondary current distribution through saline water in the above case.

The TECF has been successfully introduced in a functional prototype of personal protective equipment in the form of hand gloves. In this context commercial nitrile gloves have been modified by coating nylon and SR dots, as shown in the photograph presented in Figure 3(a). The brighter and larger dots (size ∼ 0.8-2.0 cm^2^) represent SR, while the lighter ones correspond to nylon (size ∼ 03-0.5 cm^2^). With such a design, the rubbing or tapping of the two hands generate triboelectric charges. Further the coating of the gloves in a dotted fashion allows the triboelectric charge generation even using a single hand, when the fingers and palm of the user come in contact with each other. Particularly the design presented in Figure S5 is useful for single hand operation. Figure 3(b) shows the open circuit output voltage generated out of two alternative pairs of dots of nylon and SR rubbed with fingers. Upon rubbing of the same pair of dots, an out voltage of ∼20 V across the capacitor (0.26 μF) can be realized (as shown inset) within 50 s.

**Figure 3:**
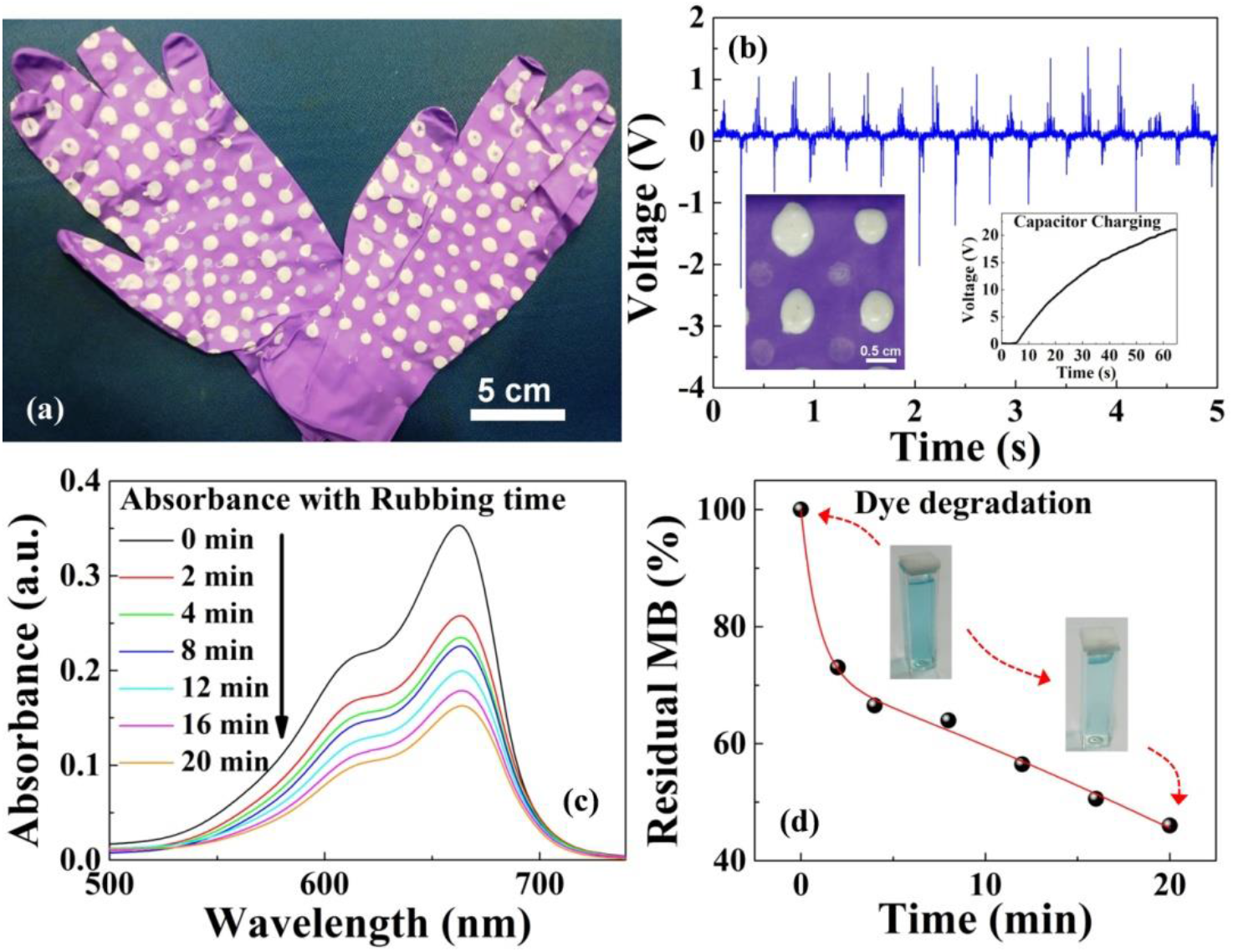
(a) A photograph of the functional prototype TECF based gloves, (b) Open circuit output voltage due to rubbing of the pair of dots (picture at the inset), while the capacitor charging by rubbing process is presented as the other inset, (c) Absorption spectra showing the degradation of methylene blue dye through rubbing of TECF based gloves, (d) The percentage degradation of methylene blue with different rubbing time, with insets presenting the pictures of the dye before and after degradation.

To realize the existence of current flow in the liquid medium, the TECF based gloves have been rubbed in the presence of a redox dye. The redox characteristics of methylene blue (MB) can be a good indicator of the existence of any current loop present in the liquid/dielectric system ^29^ and its degradation through electrochemical route can ensure the presence of a short circuit current. Figure 3(c) shows the absorption spectra of the MB dye before and after rubbing with TECF based gloves. In the absorption spectra of virgin MB (rubbing time zero), a prominent peak at ∼ 662 nm and the shoulder at ∼ 621 nm indicates the absorption due to monomer and dimer, respectively.^30^ It is noticed that the rubbing for a longer duration leads to the lowering of the absorption peak of MB indicating the degrading tendency of the dye. The degradation percentage has been extracted from the residual MB % as calculated from the absorbance values considering the relation:

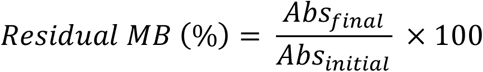

where *Abs* represents the absorption at a particular wavelength. As observed in Figure 3(d), the TECF based gloves can degrade ∼ 50% of MB dye upon 20 min of rubbing. The transformation of the coloured dye solution to a pale coloured one has also been observed by naked eye, as presented in the inset of the figure. The degradation of the redox dye is attributed to the induced current flow through the electrolyte solution on rubbing of TECF based gloves. Such situation are associated to the indirect electrolysis, where less harmful products are produced as compared to direct electrolysis process.^31^ In this case, it is expected that the active chlorine in the form of hypochlorous acid or hypochlorite ions act as the oxidizing agent to degrade MB dye. The dye degradation experiment has been further confirmed with nylon and teflon layers as presented in Figure S6 (a,b). Apart from the gradual lowering of the MB absorption peak, the development of an absorption peak of leucomethylene blue (at ∼ 268 nm) confirms the degradation of the MB solution (Figure S6 (c)). In order to ensure the role of the short circuit current, similar experiment has been performed with teflon-teflon contact system. In this case, the signature of leucomethylene blue related peak is absent in the absorption spectra, as shown in Figure S6(d). This reveals the absence of any form of hypochlorous acid since the generation of a short circuit current is not expected among teflon-teflon (same tribo-potential) pairs.

The formation of hypochlorous acid has also been substantiated through estimation and quantification of free chlorine ions using DPD (N, N-diethyl-p-phenylenediamine) colorimetric and titrimetric methods. In general, HOCl and hypochlorite ions (OCl^-^), are commonly referred as ‘free chlorine’. After rubbing the salt solution with TECF based gloves for ∼ 5 mins, DPD has been added to the salt solution. As shown in Figure 4(a), the colourless salt solution turns pink upon addition of DPD. It is well established that free chlorine can oxidize DPD to form Würster dye (which is Magenta in colour), as shown in equation presented in the Figure 4(a).

**Figure 4:**
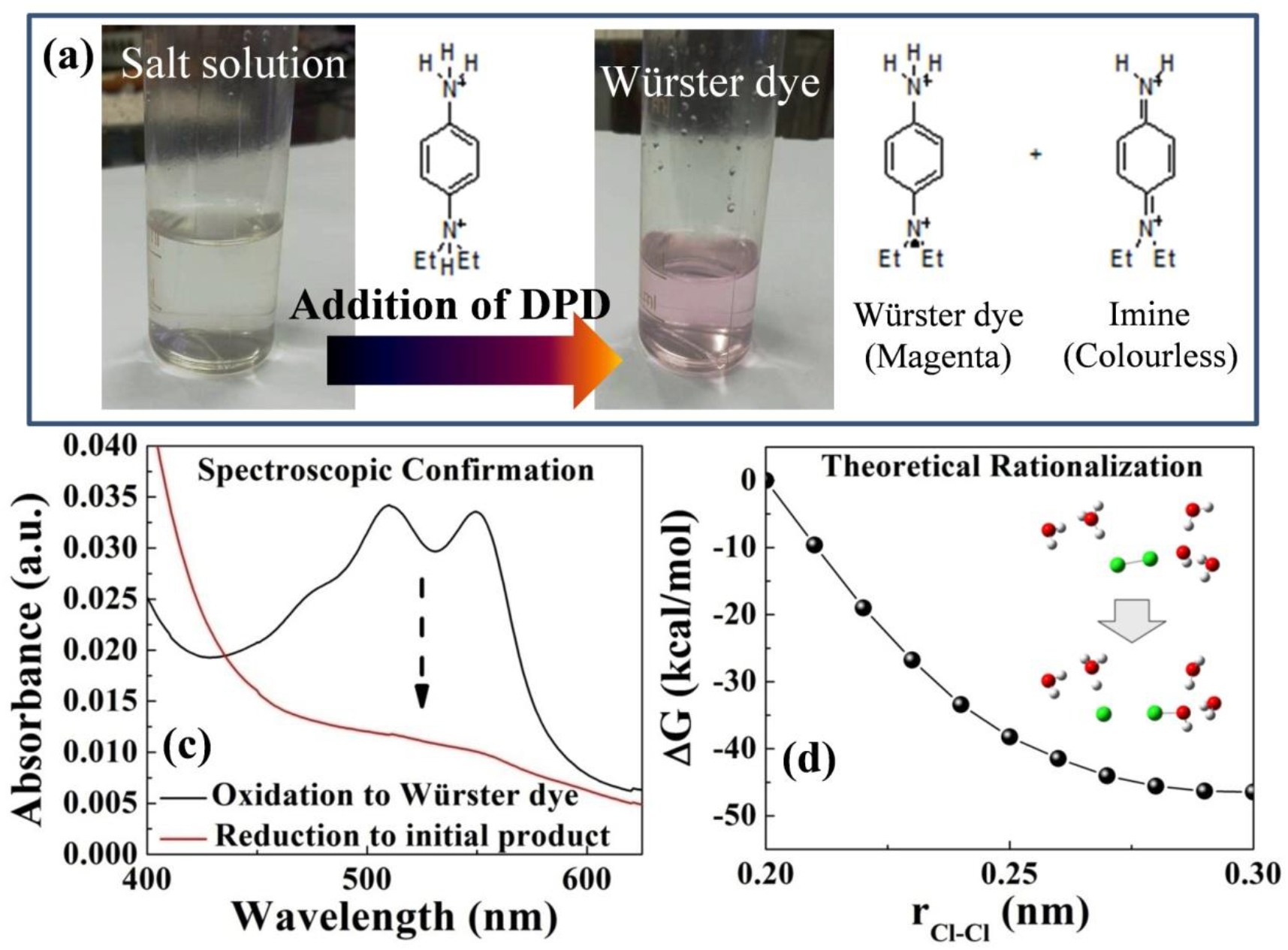
(a) Formation of Würster dye on addition of DPD into salt solution, (b) Absorbance of Würster dye and initial product, (c) Variation of total energy with inter atomic distance of Cl and O. Inset represents a scheme of HOCl formation.

The absorbance spectrum of the obtained pinkish solution (Figure 4(b)) shows the presence of a doublet peak with maxima at ∼ 511 and 551 nm, which is in accordance of the absorption peak of Würster dye.^32^ The reduction of the Würster dye using a ferrous reducing agent via titrimetric method has led to the reappearance of the initial colourless solution. The disappearance of the doublet peak in the absorbance spectrum (Figure 4(b)) indicates the regaining of the initial form of DPD. Additionally, such titrimetric analysis indicates the presence of ∼ 5 ppm of HOCl in the salt solution subjected to 5 min slow rubbing with TECF based gloves. It is worth to mention that the concentration of HOCl produced in this process is not harmful to human being even if it comes in direct contact with the skin.^33^

Next, to understand the formation of HOCl with TECF based gloves, the ongoing electrolysis process has been revisited. During the electrolysis, water takes up electrons to produce hydrogen gas and hydroxyl ions (OH^-^) at the cathode following equation (1). The OH^-^ or Cl^-^, present in the saline water gets attracted to the anode. At the anode, Cl^-^ releases an electron to form chloride radicals (Cl) and the combination of two Cl forms chlorine gas (Cl^2^), as in equation (2). Such Cl^2^ gas upon splitting from the Cl-Cl bond, can combine with water to form HOCl following equation (3).

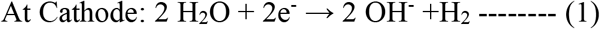

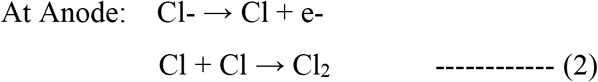

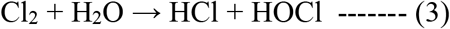

On the contrary at a lower salt concentration, OH^-^ can release electrons to the anode to form hydroxyl radicals (OH) and the combination of OH with Cl radicals can also yield HOCl, as shown in equation (4).

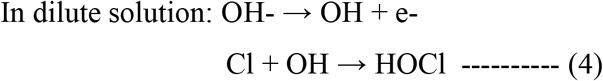

The radical reaction is extremely favourable and a barrier-less process with a free energy of - 4.2 × 10^5^ J/mole, as evident by the atomistic quantum chemical calculations. The formation of HOCl out of Cl^2^ gas in presence of OH^-^ ions on the anode surface, suggests a barrier less spontaneous process (Figure 4(d)), while the figure inset shows the scheme of formation mechanism of HOCl.

For demonstrating the antibacterial activity *P. auregionosa* has been investigated owing to its similar amino acids in the binding sites of the proteins as that of SARS-CoV-2 virus. As can be seen in Figure 5 (a-c), there is significant similarities in the amino acid sequence in lecA of *P. aeruginosa* lectin and spike RBD of SARS-CoV-2. In order to evaluate antibacterial efficacy of the developed TECF based gloves, the plate count method has been deployed. It is evident from the plate count assay (Figure 6(a,b)) that in TECF based gloves, the bacterial growth is ∼69% lower compared to control (p<0.001) and ∼64% lower compared to control TECF (nylon-nylon) based system (p<0.001). Thus it is quite evident that TECF based gloves can hinder the growth of *P. auregionosa* through their anti-microbial activity and hence can also be expected to be effective for viruses like CoV-2. In a similar fashion, the anti-microbial activity of the TECF based gloves has been examined for *E. coli*. The effectiveness of TECF can be realized from the perturbed growth of the bacteria as can be observed in the plate count assay (Figure 6(c,d)). Furthermore, in order to evaluate inhibition kinetics of the nanobiocomposites under dynamic condition as a function of time, submerged culture or growth kinetics method is adopted. Figure 6(e) clearly shows that the growth of *E. coli* subjected to the TECF is significantly retarded. The growth rate is ∼6 times lower in case of TECF compared to both usual control and TECF control (Figure 6(e)-inset) samples. Further the number of dead cells and live cells in different matrix has been evaluated by staining with DAPI and PI (Figure 6(f)). PI generally stains dead cells, whereas, DAPI stains all nuclear material irrespective of their viability. The stained image (Figure 6(f)) of control fabric and cotton fabric inoculated with bacteria shows more number of live cells than dead cells in the ratio ∼1.2. However, the number of living cells are significantly reduced in case of TECF based gloves (∼0.45). As apoptosis of cells are characterized by DNA fragmentation and consequently loss of nuclear DNA content, the ability of PI to specifically label DNA makes it useful to stain only the dead cells. The higher density of dead cells in the matrix further implies that TECF based gloves induce membrane damage, resulting into a loss of membrane potential leading to DNA fragmentation and subsequent staining of PI, thus indicating their effectiveness as an anti-microbial agent.

**Figure 5:**
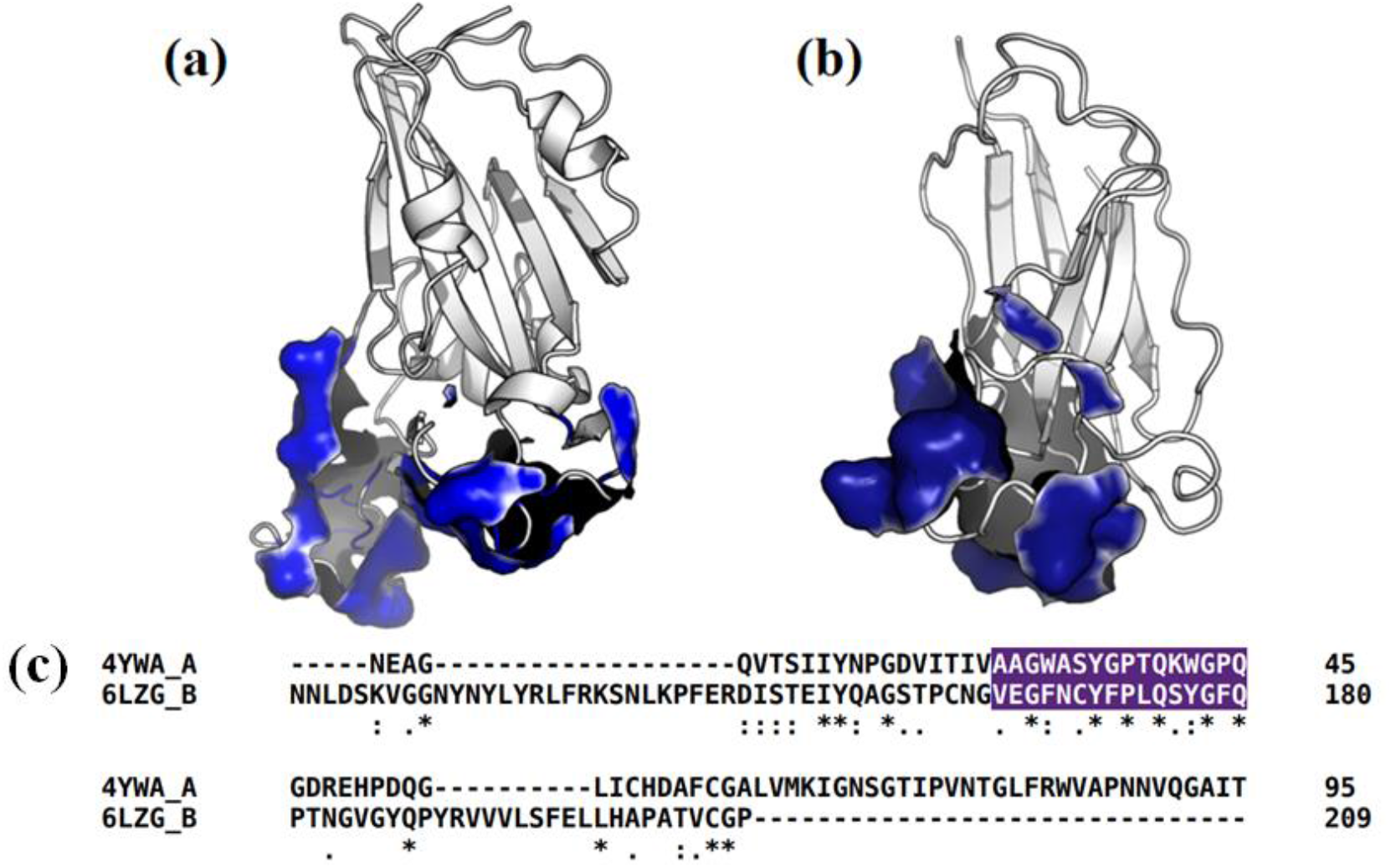
Comparison of the SARS-CoV-2 spike protein receptor binding domain (RBD) with the *P. aeruginosa* lectin lecA. (a) SARS-CoV-2 spike RBD with the binding site residues highlighted in blue. (b) *P. aeruginosa* lecA with binding site residues highlighted in blue. (c) Amino acid sequence comparison between lecA (PDB ID: 4YWA; chain A) and SARS-CoV-2 spike RBD (PDB ID: 6LZG; chain B). Binding region is highlighted in blue. Amino acids are written in one letter code; star (*) indicates identical amino acids; very similar and similar amino acids are marked with colon (:) and dot (.), respectively.

**Figure 6:**
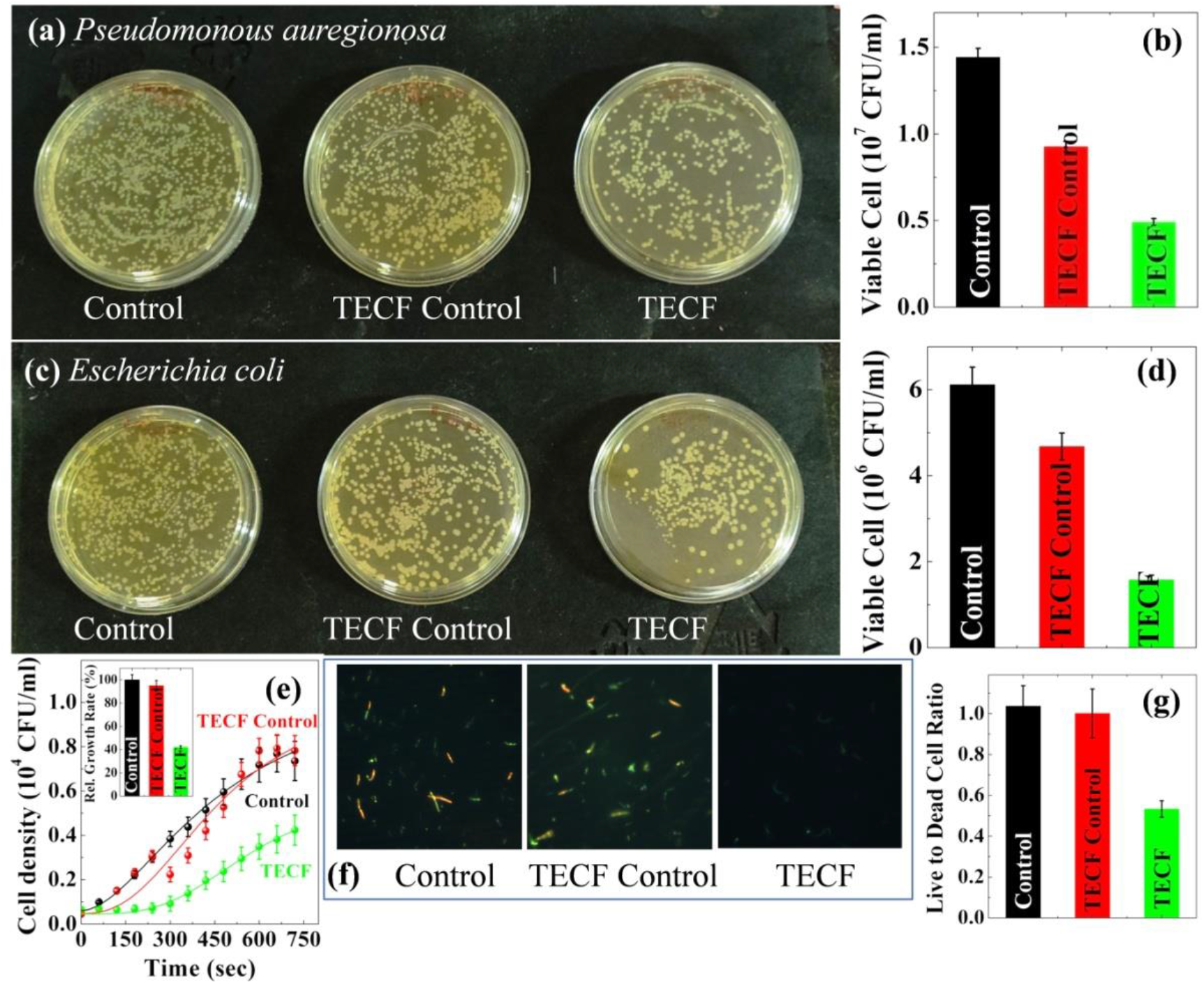
Anti-microbial activity against *P. auregionosa* and *E. coli* bacteria. (a,c) Agar-plate colony count assay, (b,d) Quantification of colonies counted in agar-plate assay. (e) Growth curve of *P. auregionosa*. Inset shows relative growth rate compared to control and TECF control samples.(f) DAPI/PI dual stained cells (*P. auregionosa)* observed under a fluorescence microscope. Blue cells are stained with DAPI, while red are stained with PI. (g) Ratio of live:dead bacterial cells as found in the DAPI/PI dual staining experiment.

The origin of the microbial activity of the TECF based gloves can be summarized in the scheme presented in Figure 7. TECF layers can be compared to a commercial battery which in presence of saline water can produce electrolyzed water. Finally to check the repeatability of hypochlorous acid formation, the TECF based gloves have been tested for number of continuous cycles for degrading the MB dye. Figure 8(a) shows the dye degradation for four continuous cycles upon rubbing the gloves. With each cycle the initial amount of dye has been added to compensate the degraded amount. The dye degradation as a result of hypochlorous acid formation is evident from decrement in the absorbance value (at ∼664 nm) throughout all the operative cycles. The consistent rate constant of the ongoing reaction (Figure 8(b)) indicates the consistent performance of the TECF based gloves. It is obvious that rubbing the gloves in presence of salt water or normal tap water (which contains dissolved salts) can trigger the formation of hypochlorous acid, which can work as a disinfectant to save us from bacteria or viruses including the SARS-COV-2 virus. The study demonstrates the self-sanitization capability of proposed TECF based gloves, making them wearable for a longer period without disposing frequently and capable of preventing the undesirable community spread of virus from otherwise uninfected frontline workers. Though the proof of concept has been demonstrated only for hand gloves, the novel TECF based wearables are potentially attractive for various other PPEs used for combating the community spread of multiple range of viral diseases.

**Figure 7:**
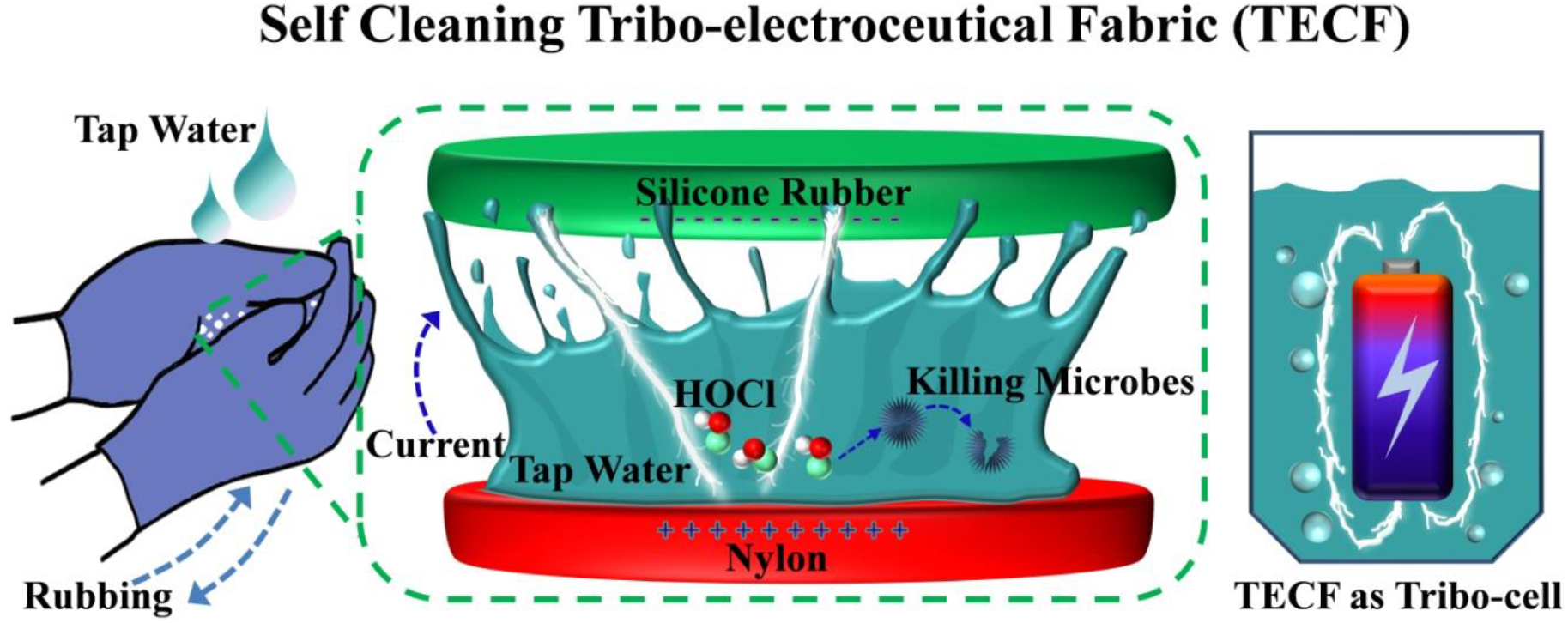
Schematic representation of self sanitization of TECF based gloves and the illustration of electrolyzed water formation by a battery.

**Figure 8:**
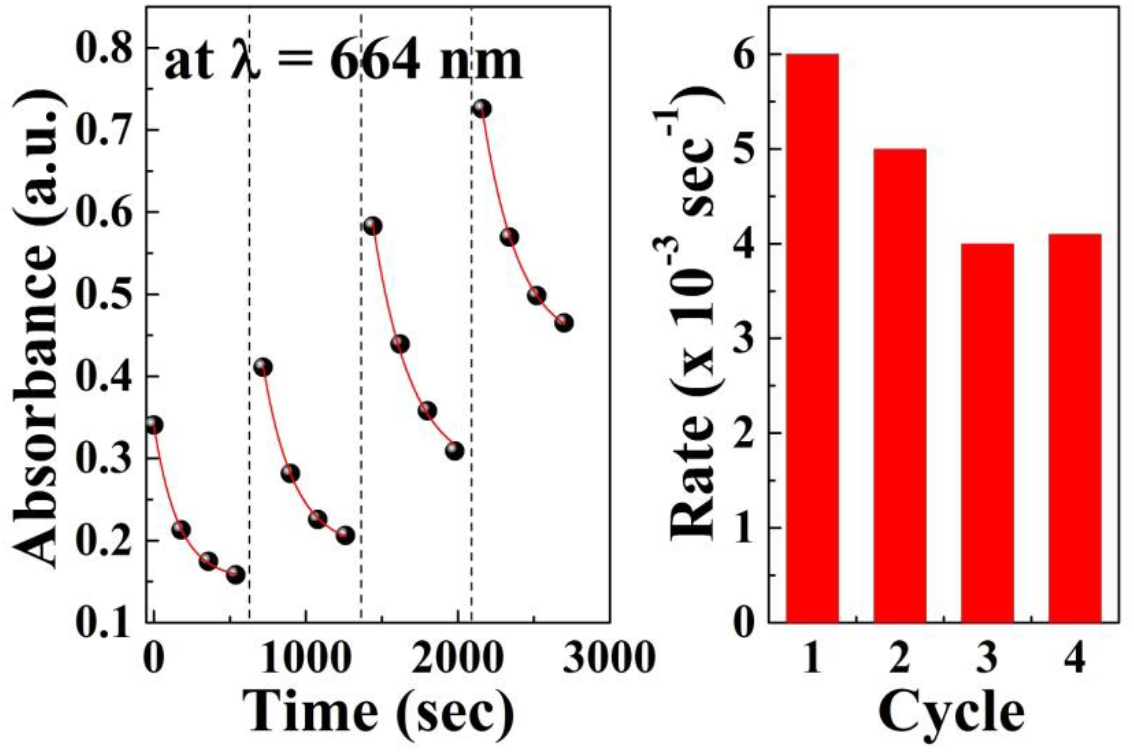
(a) MB dye degradation at different continuous cycles, (b) rate constant of the dye degradation reaction.

## Conclusions

We report on the fabrication of tribo-electroceutical fabric (TECF) based hand gloves with self sanitization ability. The triboelectric charges from Silicone Rubber (SR)-Nylon pair has been deployed to activate the self sanitization process. The power generation ability of the SR-Nylon pair has been understood upon investigating the contact electrification process. The TECF pair can generate as high as 20 V under and a maximum power of ∼41 μW/cm^2^ with a capability to charge a 0.26 F capacitor up to 65 V within 50 sec. Further generation of a short circuit current has been witnessed when the TECF pair is rubbed in presence of saline water. In this investigation, TECF pair based functional prototype gloves have been developed on commercially available nitrile platform. The evolution of short circuit current and subsequent formation of hypochlorous acid through the rubbing of such prototype gloves has been realized. Further theoretical modelling has predicted that the event of TECF led formation of hypochlorous acid is energetically favourable and a barrier-less process. It has been found that rubbing of such TECF based gloves for 5 min can generate ∼ 5 ppm of hypochlorous acid. The self-sanitization ability of the TECF based gloves has been demonstrated through antibacterial activities against bacteria like *P. auregionosa* and *E. Coli*. The demonstration of excellent antibacterial activity against *P. auregionosa* suggests the potential application of the TECF based gloves against the ongoing battle against coronavirus. In addition to the use in common PPE, the developed method is expected to find relevance in sanitization of various surfaces (e.g. table, chairs etc.) without using any antibacterial chemical.

## Acknowledgements

SB acknowledges CSIR for financial assistance under SRA (Scientists Pool Scheme). SKP acknowledges INAE for Abdul Kalam Technology Innovation National Fellowship. TSD acknowledges the funding by J. C. Bose Fellowship.

## Supporting Information

Polarity based electrical output generation; mechanism of contact electrification; output power and voltage generation across different resistances; Capacitor charging through rubbing/sliding of Teflon-Nylon; Additional design of SR and Nylon coated Nitrile gloves; dye degradation study through rubbing of Teflon-Nylon layers.

