## Supplementary Information for "Development of Tribo-Electroceutical Fabrics for Potential Application in Self-sanitizing Personal Protective Equipment (PPE)"

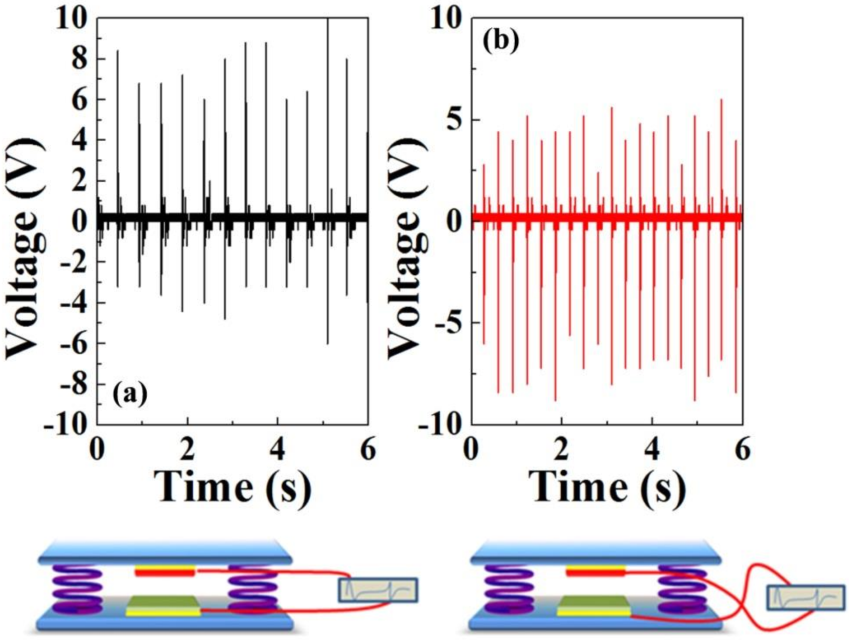


**Figure S1**: Output generation through (a) Forward and (b) Reverse connection

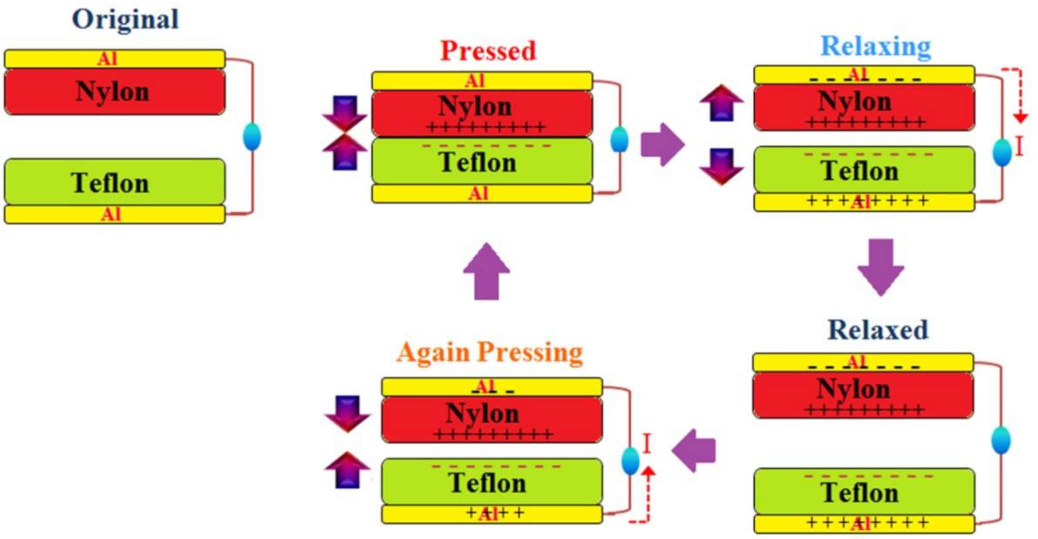


**Figure S2**: Mechanism of contact electrification.

In brief, it is known that Teflon and Nylon are negative and positive charge seeking materials respectively. The contact between the two layers leads to the charge transfer, while the subsequent separation of these layers leads to the development of potential. Further the re-contact between the two leads to the development of an inverted potential which is reflected as output voltage with opposite polarity.

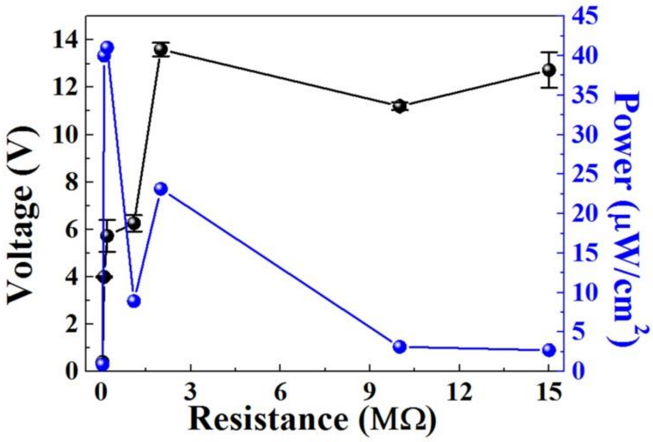


**Figure S3**: Variation of output power and voltage across different resistances.

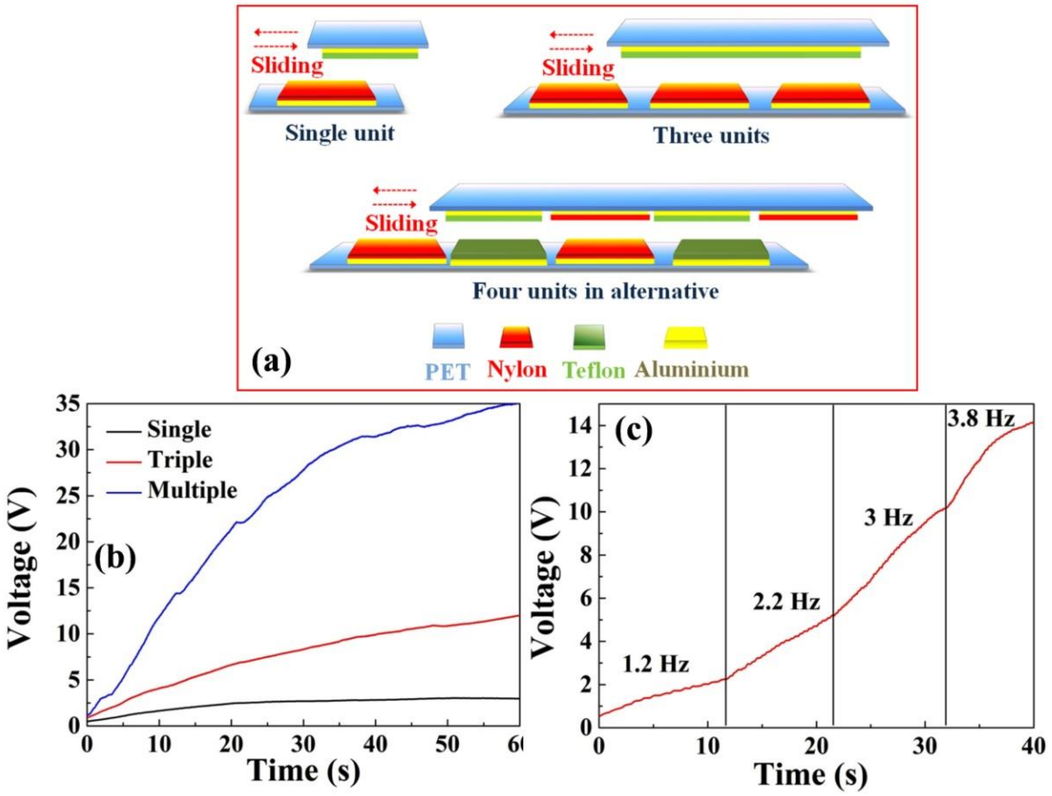


**Figure S4**: (a) Scheme of different units of Teflon-Nylon for contact electrification through rubbing/sliding, (b) Capacitor charging with different units. (c) Capacitor charging with different frequencies of sliding.

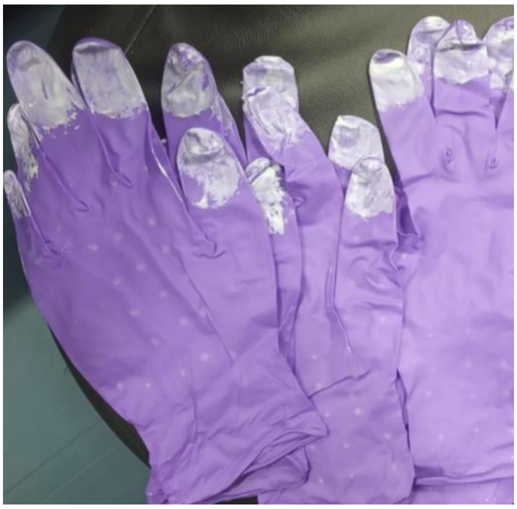

**Figure S5:** An unique design of SR and Nylon coated Nitrile gloves. SR is coated on the tip of the fingers, while Nylon is deposited on the palm.

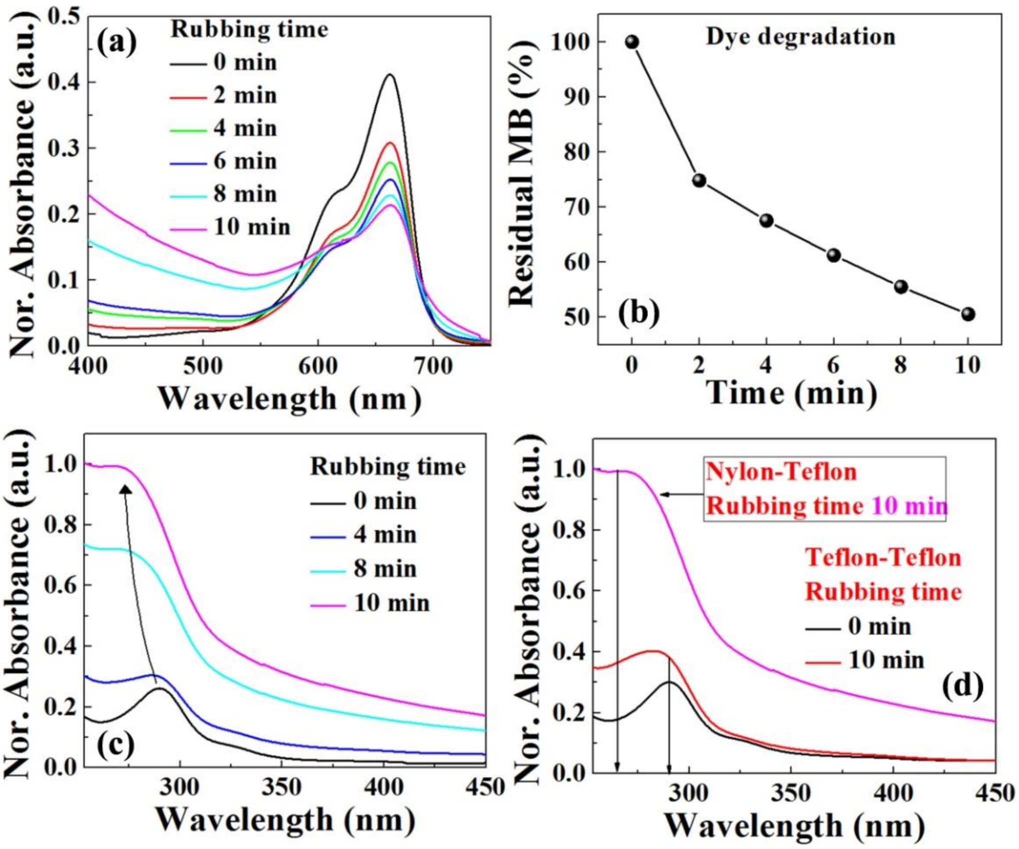


**Figure S6:** (a) Methylene blue degradation through rubbing of Teflon-Nylon layers, (b) Degradation percentage with different time of rubbing, (c) Formation of leucomethylene blue (at 265 nm) through rubbing of Teflon-Nylon layers, (d) Comparison of the formation of leucomethylene blue for Teflon-Nylon and Teflon-Teflon layers.
